## Supplementary Material for "Reconstructing the phylogeny and evolutionary history of freshwater fishes (Nemacheilidae) across Eurasia since early Eocene"

**Table S1.**

List of analysed samples, their identification, geographical origin, voucher number and GenBank accession numbers for their sequences. Voucher numbers starting with 'A' refer to the collection of IAPG, Liběchov, Czech Republic; 'CMK' numbers refer to the collection of Maurice Kottelat; 'GenBank' refers to sequences from GenBank; 'ZRC' to samples housed in the Lee Kong Chian Natural History Museum, National University of Singapore, Singapore.

| Species name | Country | Province | River drainage | voucher | Cyt b | RAG 1 | IRBP2 | MYH6 | RH 1 | EGR 3 |
|---|---|---|---|---|---|---|---|---|---|---|
| **EASTERN CLADE** | | | | | | | | | | |
| ***Karstsinnectes acridorsalis*** | China | Guangxi | Pearl | GenBank | ON116515 | OP473644 | - | OP473686 | OP473776 | OP473850 |
| ***Karstsinnectes anophthalmus*** | China | Guangxi | Pearl | GenBank | ON116506 | OP473637 | OP473907 | OP473664 | OP473763 | OP473828 |
| ***Karstsinnectes parvus*** | China | Guangxi | Pearl | GenBank | ON116520 | OP473651 | OP473900 | OP473693 | - | OP473859 |
| ***Lefua costata*** | Korea | Gangwon | Cheon Jin Cheon | A1895<br>A6942 | PP279919<br>KP738591 | PP315753<br>KP738551 | PP280130<br>KP738511 | PP280341<br>OL191348 | -<br>- | -<br>PP259693 |
| ***Lefua torrentis*** | Japan | Hyogo | Kako | A1050<br>A1051<br>A1052 | PP279878<br>PP279879<br>PP279880 | PP315711<br>PP315712<br>PP315714 | PP280091<br>PP280092<br>PP280093 | PP280288<br>PP280289<br>PP280291 | PP259745<br>PP259747<br>PP259748 | -<br>-<br>- |
| ***Micronemacheilus bailianensis*** | China | Guangxi | Pearl | GenBank | ON116504 | OP473620 | OP473876 | OP473662 | OP473759 | OP473825 |
| ***Micronemacheilus cruciatus*** | Vietnam | Thua Tien-Hue | Song Bu Lu | A3293<br>A3294 | PP279965<br>PP279966 | PP315801<br>PP315802 | -<br>- | PP280394<br>PP280395 | PP259847<br>PP259848 | PP259670<br>PP259671 |
| ***Micronemacheilus longibarbatus*** | China | Guangxi | Pearl | GenBank | ON116508 | OP473625 | OP473879 | OP473666 | OP473761 | OP473833 |
| ***Micronemacheilus pulcherimus*** | China | Guangxi | Pearl | A8690<br>A8691 | PP280043<br>PP280044 | PP315878<br>PP315879 | PP280229<br>PP280230 | PP280508<br>PP280509 | PP259968<br>PP259969 | PP259700<br>- |
| ***Oreonectes guananensis*** | China | Guangxi | Pearl | GenBank | ON116507 | OP473623 | OP473878 | OP473665 | OP473750 | OP473830 |
| ***Oreonectes luochengensis*** | China | Guangxi | Pearl | GenBank | ON116495 | - | OP473882 | OP473670 | OP473749 | OP473836 |
| ***Oreonectes platycephalus*** | China | Hong Kong | Tai Tam | A1674<br>A1675<br>A1676 | PP279902<br>PP279903<br>PP279904 | PP315737<br>PP315738<br>PP315739 | PP280115<br>PP280116<br>PP280117 | PP280323<br>PP280324<br>PP280325 | PP259777<br>PP259778<br>PP259779 | -<br>-<br>- |

| | | | | | | | | | | |
|---|---|---|---|---|---|---|---|---|---|---|
| | | | | GenBank | ON116528 | OP473652 | OP473904 | OP473696 | OP473745 | OP473862 |
| ***Oreonectes* cf. *platycephalus 1*** | China | Guangxi | Pearl | A9065 | PP280053 | PP315888 | PP280239 | PP280518 | PP259977 | PP259707 |
| ***Oreonectes* cf. *platycephalus 2*** | China | Guangxi | Pearl | A8697 | PP280045 | PP315880 | PP280231 | PP280510 | PP259970 | PP259701 |
| ***Oreonectes* cf. *platycephalus 3*** | Vietnam | Lang Son | Pearl | A10603 | PP279881 | - | - | PP280292 | - | - |
| ***Oreonectes polystigmus*** | China | Guangxi | Pearl | GenBank | ON116514 | OP473614 | OP473871 | OP473657 | OP473746 | OP473856 |
| ***Oreonectes* sp.** | China | Guangxi | Pearl | A8963 | PP280047 | PP315882 | PP280233 | PP280512 | PP259972 | PP259702 |
| | | | | A8964 | PP280048 | PP315883 | PP280234 | PP280513 | - | PP259703 |
| ***Paranemachilus genilepis*** | China | Guangxi | Pearl | GenBank | ON116497 | OP473630 | OP473885 | OP473673 | OP473752 | OP473839 |
| ***Paranemachilus pingguoensis*** | China | Guangxi | Pearl | GenBank | ON116500 | OP473634 | OP473888 | OP473676 | OP473755 | OP473843 |
| ***Paranemachilus zhengbaoshani*** | China | Guangxi | Pearl | A9153 | PP280054 | PP315889 | PP280240 | PP280519 | PP259978 | PP259708 |
| ***Sundoreonectes sabanus*** | Malaysia | Sabah | Baram | A1844 | PP279914 | PP315748 | PP280126 | PP280334 | PP259789 | PP259651 |
| | | | | A1845 | PP279915 | PP315749 | PP280127 | PP280335 | PP259790 | PP259652 |
| ***Traccatichthys pulcher*** | China | Guangxi | Pearl | A1804 | PP279910 | PP315744 | PP280122 | PP280330 | PP259785 | - |
| | | | | A1805 | PP279911 | PP315745 | PP280123 | PP280331 | PP259786 | - |
| | | | | A8681 | PP280042 | PP315877 | PP280228 | PP280507 | PP259967 | PP259699 |
| ***Traccatichthys taeniatus*** | Vietnam | Nghe An | Lam | A3175 | PP279960 | PP315796 | PP280167 | PP280387 | PP259840 | PP259666 |
| | | | | A3176 | PP279961 | PP315797 | PP280168 | PP280388 | PP259841 | - |
| | Laos | Houaphan | Lam | CMK 25931 | PP279871 | PP315710 | PP280088 | PP280249 | PP259738 | - |
| ***Traccatichthys* cf. *taeniatus*** | Vietnam | Quang Nam | Cau Do | A9575 | PP280060 | - | - | - | - | - |
| ***Traccatichthys zispi*** | China | Hainan | - | GenBank | ON116518 | OP473648 | OP473898 | OP473691 | OP473780 | OP473861 |
| ***Troglonectes barbatus*** | China | Guizhou | Pearl | GenBank | ON116501 | OP473635 | OP473889 | OP473678 | OP473756 | - |
| ***Troglonectes daqikongensis*** | China | Guizhou | Pearl | GenBank | ON116526 | OP473641 | - | OP473683 | OP473773 | OP473849 |
| ***Troglonectes dongganensis*** | China | Guangxi | Pearl | GenBank | ON116503 | OP473617 | OP473875 | OP473661 | OP473757 | OP473847 |
| ***Troglonectes donglanensis*** | China | Guangxi | Pearl | GenBank | ON116505 | OP473621 | OP473877 | OP473663 | OP473762 | OP473826 |
| ***Troglonectes duanensis*** | China | Guangxi | Pearl | GenBank | ON116509 | OP473622 | OP473880 | OP473667 | OP473764 | OP473831 |

| | | | | | | | | | | |
|---|---|---|---|---|---|---|---|---|---|---|
| ***Troglonectes elongatus*** | China | Guangxi | Pearl | GenBank | ON116502 | OP473616 | OP473874 | OP473660 | OP473758 | OP473848 |
| ***Troglonectes furcocaudalis*** | China | Guangxi | Pearl | GenBank | ON116512 | OP473628 | OP473883 | OP473671 | OP473767 | OP473837 |
| ***Troglonectes jiarongensis*** | China | Guizhou | Pearl | GenBank | ON116527 | OP473643 | OP473894 | OP473685 | OP473769 | OP473846 |
| ***Troglonectes lihuensis*** | China | Guangxi | Pearl | GenBank | ON148332 | OP473618 | OP473892 | OP473688 | - | OP473852 |
| ***Troglonectes macrolepis*** | China | Guangxi | Pearl | GenBank | ON116498 | OP473632 | OP473886 | OP473674 | OP473753 | OP473841 |
| ***Troglonectes microphthalmus*** | China | Guangxi | Pearl | A8988<br>A8989<br>GenBank | PP280049<br>PP280050<br>ON116494 | PP315884<br>PP315885<br>OP473631 | PP280235<br>PP280236<br>OP473872 | PP280514<br>PP280515<br>OP473659 | PP259973<br>PP259974<br>OP473748 | PP259704<br>PP259705<br>OP473840 |
| ***Troglonectes retrodorsalis*** | China | Guangxi | Pearl | GenBank | ON116511 | OP473627 | OP473873 | OP473669 | OP473766 | OP473835 |
| ***Troglonectes shuilongensis*** | China | Guizhou | Pearl | GenBank | ON116522 | OP473636 | OP473891 | OP473679 | OP473768 | OP473834 |
| ***Troglonectes translucens*** | China | Guangxi | Pearl | GenBank | ON116510 | OP473626 | OP473881 | OP473668 | OP473765 | OP473832 |
| ***Yunnanilus pleurotaenia*** | China | Yunnan | Yangtze | A2967<br>A2968<br>A2969 | PP279954<br>PP279955<br>PP279956 | PP315791<br>PP315792<br>PP315793 | PP280163<br>PP280164<br>PP280165 | PP280381<br>PP280382<br>PP280383 | PP259836<br>PP259837<br>PP259838 | PP259664<br>PP259665<br>- |
| **NORTHERN CLADE** | | | | | | | | | | |
| ***Barbatula*** | France | Haute- | Garonne | CMK18464 | PP279842 | PP315677 | - | PP280251 | - | - |
| ***barbatula*** | | Garonne | | _1CMK184 | PP279843 | PP315678 | - | PP280252 | PP259711 | PP259624 |
| | Russia | Unknown | unknown | 64_2 | PP279979 | PP315816 | PP280179 | PP280409 | PP259858 | - |
| | | | | A4013 | PP279980 | PP315817 | PP280180 | PP280410 | - | PP259674 |
| | Germany | Nordrhein- | Rhine | A4015 | PP279922 | PP315756 | PP280133 | PP280344 | PP259798 | PP259655 |
| | | Westfalen | | A2046 | PP279923 | PP315757 | PP280134 | PP280345 | PP259799 | - |
| | Czechia | Liberecky | Elbe | A2047 | PP279923 | PP315777 | PP280150 | PP280367 | PP259822 | - |
| | | | | A2587 | PP279941 | PP315778 | PP280151 | PP280368 | PP259823 | - |
| | Czechia | Stredocesky | Elbe | A2588 | KP738604 | KP738564 | KP738524 | PP280499 | PP259955 | PP259697 |
| | | | | A8393 | KP738605 | KP738565 | KP738525 | OL191359 | PP259956 | - |
| | Poland | Dolnośląskie | Oder | A8394 | PP279952 | PP315789 | PP280161 | PP280379 | PP259834 | - |
| | | | | A2957 | PP279953 | PP315790 | PP280162 | PP280380 | PP259835 | - |
| | | | | A2958 | | | | | | |

| | | | | | | | | | | |
|---|---|---|---|---|---|---|---|---|---|---|
| ***Barbatula* cf. *compressirostris*** | Mongolia<br><br>Mongolia | Khovd<br><br>Khovd | Khovd Tsendkhar<br>Khovd | CMK19564_1<br>CMK19564_2<br>CMK19590_1CMK19590_2 | PP279848<br>PP279849<br>PP279852<br>PP279853 | PP315683<br>PP315684<br>PP315687<br>PP315688 | PP280067<br>PP280068<br>PP280071<br>PP280072 | PP280257<br>PP280258<br>PP280261<br>PP280262 | PP259716<br>PP259717<br>PP259720<br>PP259721 | PP259626<br>-<br>PP259628<br>- |
| ***Barbatula dgebuadzei*** | Mongolia | Bayankhongor | Baydrag | CMK19594_1<br>CMK19594_2 | PP279854<br>PP279855 | PP315689<br>PP315690 | PP280073<br>PP280074 | PP280263<br>PP280264 | PP259722<br>PP259723 | PP259629<br>- |
| ***Barbatula karabanowi*** | Mongolia | Khovd | Bulgan Gol | CMK19584_1<br>CMK19584_2 | PP279850<br>PP279851 | PP315685<br>PP315686 | PP280069<br>PP280070 | PP280259<br>PP280260 | PP259718<br>PP259719 | PP259627<br>- |
| ***Barbatula oreas*** | Japan | Hokkaido | Shinkawa | A11223<br>A11224 | PP279887<br>PP279888 | PP315722<br>PP315723 | PP280097<br>PP280098 | PP280300<br>PP280301 | PP259754<br>PP259755 | PP259638<br>- |
| ***Barbatula* sp. Korea** | Korea | Gangwon | Cheon Jin Cheon | A1888 | PP279918 | PP315752 | - | PP280340 | - | - |
| ***Barbatula* sp. Tuul** | Mongolia | Ulaanbaatar | Selenga | A4246<br>A4247<br>A4248<br>A4250 | PP279981<br>PP279982<br>PP279983<br>PP279985 | -<br>PP315818<br>PP315819<br>PP315821 | -<br>PP280181<br>PP280182<br>PP280184 | PP280412<br>PP280413<br>PP280414<br>PP280416 | PP259861<br>-<br>PP259862<br>PP259863 | PP259676<br>-<br>-<br>- |
| ***Barbatula toni*** | Russia | Primorsky | Amur | A3171 | PP279959 | - | - | PP280386 | - | - |
| ***Barbatula* cf. *toni*** | Mongolia<br><br>Mongolia | Khövgöl<br><br>Ulaanbaatar | Selenga<br><br>Selenga | CMK19540_1<br>CMK19540_2A4249 | PP279844<br>PP279845<br>PP279984 | PP315679<br>PP315680<br>PP315820 | PP280063<br>PP280064<br>PP280183 | PP280253<br>PP280254<br>PP280415 | PP259712<br>PP259713<br>- | -<br>-<br>PP259677 |
| ***Claea dabryi*** | China | Sichuan | Yangtze | GenBank | MG238214 | MG237922 | MG238312 | - | MG238015 | - |
| ***Triplophysa baotianensis*** | China | Guizhou | Pearl | GenBank | MT992550 | OP473612 | OP473868 | OP473655 | - | OP473864 |
| ***Triplophysa bleekeri*** | China | no detail | Yangtze | GenBank | MG238298<br>KX373847 | MG238003<br>MG725561 | MG238415<br>MG698830 | -<br>MG698529 | -<br>MG697779 | -<br>- |
| ***Triplophysa brevicauda*** | China | no detail | Yangtze | GenBank | MG238300 | MG238005 | MG238417 | - | MG238107 | - |
| ***Triplophysa dalaica*** | China | Gansu | no details | GenBank | MG697586 | - | MG698831 | MG698530 | MG697806 | - |
| ***Triplophysa dorsalis*** | Kazachstan | Almaty | Lake Balkash | A5377<br>A5378<br>A5380 | PP280001<br>PP280002<br>PP280004 | PP315838<br>PP315839<br>PP315841 | PP280191<br>PP280192<br>PP280194 | PP280444<br>PP280445<br>PP280447 | PP259893<br>PP259894<br>PP259895 | PP259684<br>PP259685<br>- |

| | | | | | | | | | | |
|---|---|---|---|---|---|---|---|---|---|---|
| | | | | A5383 | PP280007 | PP315844 | PP280196 | PP280450 | PP259898 | - |
| ***Triplophysa grahami*** | China | Yunnan | Yangtze | A1663<br>A2933 | MK608125<br>PP279948 | OL191414<br>PP315785 | MT536722<br>- | OL191277<br>PP280375 | PP259776<br>PP259830 | PP259648<br>- |
| ***Triplophysa gundriseri*** | Mongolia | Khövgöl | Tes Gol | CMK19543_1<br>CMK19543_2 | PP279846<br>PP279847 | PP315681<br>PP315682 | PP280065<br>PP280066 | PP280255<br>PP280256 | PP259714<br>PP259715 | PP259625<br>- |
| ***Triplophysa huapingensis*** | China | Guangxi | Pearl | GenBank | MG697589 | - | MG698834 | MG698537 | MG697870 | - |
| ***Trplophysa labiata*** | Kazachstan | Almaty | Lake Balkash | A5381<br>A5384<br>A5385 | PP280005<br>PP280008<br>PP280009 | PP315842<br>PP315845<br>PP315846 | -<br>PP280197<br>PP280198 | PP280448<br>PP280451<br>PP280452 | PP259896<br>PP259899<br>PP259900 | -<br>-<br>- |
| ***Triplophysa leptosoma*** | China | no detail | Yangtze | GenBank | KX373839 | - | MG698825 | MG698524 | MG697601 | - |
| ***Triplophysa luochengensis*** | China | Guangxi | Pearl | A8990<br>A8991 | PP280051<br>PP280052 | PP315886<br>PP315887 | PP280237<br>PP280238 | PP280516<br>PP280517 | PP259975<br>PP259976 | PP259706<br>- |
| ***Triplophysa nandanensis*** | China | Guangxi | Pearl | GenBank | MG697588 | - | MG698833 | MG698536 | MG697869 | - |
| ***Triplophysa nanpanjiangensis*** | China | Yunnan | Pearl | GenBank | MG238302 | MG238007 | MG238419 | - | MG238109 | - |
| ***Triplophysa nasobarbatula*** | China | Guizhou | Pearl | GenBank | ON116529 | OP473653 | OP473869 | OP473697 | OP473781 | OP473865 |
| ***Triplophysa obscura*** | China | no detail | Yangtze | GenBank | MG238304 | MG238009 | MG238421 | - | MG238111 | - |
| ***Triplophysa orientalis*** | China | Qinghai | Yangtze | GenBank | KX373846 | - | MG698829 | MG698528 | MG697755 | - |
| ***Triplophysa pseudoscleroptera*** | China | Qinghai | Yangtze | GenBank | MG697585 | - | MG698828 | MG698527 | MG697684 | - |
| ***Triplophysa rosa*** | China | Wulong | Yangtze | GenBank | MG697587 | MG725565 | MG698832 | MG698535 | MG697868 | - |
| ***Triplophysa scleroptera*** | China | No details | Yangtze | GenBank | MG238307<br>KX373833 | MG238012<br>MG725554 | MG238424<br>MG698838 | <br>MG698534 | MG238113<br>MG697908 | -<br>- |
| ***Triplophysa siluroides*** | China | no details | no details | 1797 | MT536720 | EF063156 | MT536723 | OL191278 | PP259784 | PP259650 |
| ***Triplophysa stenura*** | China | Yunnan | Yangtze | 2935<br>2938<br>2939<br>GenBank | PP279949<br>PP279950<br>PP279951<br>MG697583 | PP315786<br>PP315787<br>PP315788<br>MG725550 | PP280158<br>PP280159<br>PP280160<br>MG698824 | PP280376<br>PP280377<br>PP280378<br>MG698523 | PP259831<br>PP259832<br>PP259833<br>MG697592 | -<br>PP259662<br>PP259663<br>- |
| ***Triplophysa stolickai*** | China | no detail | no details | GenBank | MG697582 | MG725535 | MG698822 | MG698521 | MG697590 | - |

| | | | | | | | | | | |
|---|---|---|---|---|---|---|---|---|---|---|
| ***Triplophysa strauchi*** | Kazachstan | Almaty | Lake Balkash | 5367 | PP280000 | - | - | - | - | - |
| | | | | 5379 | PP280003 | PP315840 | PP280193 | PP280446 | - | PP259686 |
| | | | Syr Darya | 5382 | PP280006 | PP315843 | PP280195 | PP280449 | PP259897 | - |
| | Kyrgyzstan | Naryn | | 11496 | MT536721 | OL191500 | MT536724 | OL191382 | PP259757 | PP259639 |
| | | | | 11497 | PP279889 | PP315725 | PP280100 | PP280303 | PP259758 | - |
| ***Triplophysa tenuis*** | China | no detail | no details | GenBank | MG697584 | MG725556 | MG698827 | MG698526 | MG697626 | - |
| ***Triplophysa wuweiensis*** | China | Gansu | no details | GenBank | KX373838 | - | MG698823 | MG698522 | MG697591 | - |
| **INDOCHINESE CLADE** | | | | | | | | | | |
| ***Homatula anguillioides*** | China | Yunnan | Mekong | GenBank | HM010583 | HM010669 | MG238315 | - | MG238018 | - |
| ***Homatula change*** | China | Yunnan | Mekong | GenBank | - | - | MG238318 | - | MG238021 | - |
| ***Homatula cryptoclathrata*** | China | Yunnan | Salween | GenBank | HM010569 | HM010663 | MG238332 | - | - | - |
| ***Homatula disparizona*** | China | Yunnan | Red | GenBank | MG238218 | MG237926 | MG238321 | - | MG238023 | - |
| ***Homatula laxiclathra*** | China | Shanxi | Yellow | GenBank | MG238219 | MG237927 | MG238322 | - | MG238024 | - |
| ***Homatula longidorsalis*** | China | Yunnan | Pearl | GenBank | HM010522 | HM010618 | MG238324 | - | MG238026 | - |
| ***Homatula potanini*** | China | Sichuan | Yangtze | A1788 | PP279908 | PP315742 | PP280120 | PP280328 | PP259782 | PP259649 |
| | | | | A1789 | PP279909 | PP315743 | PP280121 | PP280329 | PP259783 | - |
| ***Homatula pycnolepis*** | China | Yunnan | Mekong | A2973 | PP279957 | PP315794 | PP280166 | PP280384 | PP259839 | - |
| ***Homatula variegata*** | China | Sichuan | Yangtze | A1459 | PP279900 | PP315735 | PP280112 | PP280319 | PP259773 | PP259646 |
| | | | | A1460 | PP279901 | PP315736 | PP280113 | PP280320 | PP259774 | - |
| ***Homatula wuliangensis*** | China | Yunnan | Mekong | GenBank | HM010517 | HM010609 | MG238336 | - | MG238036 | - |
| ***'Nemacheilus' arenicolus*** | Laos | Bolikhamsay | Mekong | CMK21208 | | | | PP280265 | PP259724 | - |
| | | | | _2 | | | | PP280266 | PP259725 | PP259630 |
| | | | | CMK21208 | MW512964 | MW513090 | MW513213 | | | |
| | | | | _3 | MW512965 | MW513091 | MW513214 | | | |
| ***'Nemacheilus' argyrogaster*** | Laos | Sekong | Mekong | CMK21521 | | | | PP280267 | PP259726 | PP259631 |
| | | | | _1 | | | | PP280268 | PP259727 | - |
| | | | | CMK21521 | MW512988 | MW513111 | MW513237 | | | |
| | | | | _2 | MW512989 | MW513112 | MW513238 | | | |

| | | | | | | | | | | |
|---|---|---|---|---|---|---|---|---|---|---|
| ***'Nemacheilus' banar*** | Vietnam | Kontum | Mekong | A3307 | PP279967 | PP315803 | PP280172 | PP280396 | PP259849 | PP259672 |
| | | | | A3308 | MW512968 | MW513094 | MW513217 | PP280397 | PP259850 | - |
| ***'Nemacheilus' cleopatra*** | Vietnam | Gia Lai | Song Ba | A3256 | MW512982 | MW513106 | MW513231 | PP280392 | PP259845 | PP259669 |
| | | | | A3257 | MW512983 | MW513107 | MW513232 | PP280393 | PP259846 | - |
| ***Rhyacoschistura suber*** | Laos | Xiengkhuang | Mekong | CMK22643 | PP279857 | PP315692 | - | PP280270 | PP259729 | PP259632 |
| | Laos | Xaysomboon | Mekong | CMK22484 | PP279856 | PP315691 | PP280079 | PP280269 | PP259728 | - |
| ***Schistura amplizona*** | China | Yunnan | Mekong | GenBank | MG238243 | MG237949 | MG238357 | - | MG238056 | - |
| ***Schistura bolavensis*** | Laos | Champasak | Mekong | A4618 | KP738575 | KP738535 | KP738495 | OL191315 | PP259869 | PP259681 |
| | | | | A4620 | KP738576 | KP738536 | KP738496 | PP280421 | PP259870 | - |
| ***Schistura bucculenta*** | China | Yunnan | Mekong | GenBank | JN837654 | JN837666 | - | - | - | - |
| ***Schistura callichroma*** | China | Yunnan | Red | GenBank | MG238244 | MG237950 | MG238359 | - | MG238057 | - |
| ***Schistura caudofurca*** | China | Yunnan | Red | GenBank | MG238245 | MG237951 | MG238360 | - | MG238059 | - |
| ***Schistura* cf. *amplizona*** | Thailand | Loei | Mekong | A2375 | PP279932 | PP315766 | PP280144 | PP280355 | PP259808 | PP259658 |
| ***Schistura* cf. *palma*** | Thailand | Loei | Mekong | 2376 | PP279933 | PP315767 | - | PP280356 | PP259809 | PP259659 |
| ***Schistura* cf. *schultzei*** | Thailand | Loei | Mekong | 2522 | PP279934 | PP315768 | PP280145 | PP280359 | PP259813 | PP259660 |
| | | | | 2523 | PP279935 | PP315769 | PP280146 | PP280360 | PP259814 | - |
| ***Schistura cryptofasciata*** | China | Yunnan | Salween | GenBank | MG238250 | MG237956 | MG238366 | - | MG238063 | - |
| ***Schistura desmotes*** | Thailand | Chiang Mai | Chao Phraya | A1180 | PP279892 | PP315728 | - | PP280306 | PP259761 | PP259640 |
| | | | | A1181 | PP279893 | PP315729 | - | PP280307 | PP259762 | - |
| | | | | A1182 | PP279894 | PP315730 | - | PP280308 | PP259763 | - |
| ***Schistura dubia*** | Thailand | Phrae | Chao Phraya | GenBank | MK301364 | - | - | - | - | - |
| ***Schistura fasciolata*** | China | Guangxi | Pearl | A5300 | KP738579 | KP738539 | KP738499 | PP280441 | PP259888 | PP259683 |
| | | | | A5301 | KP738580 | KP738540 | KP738500 | PP280442 | PP259889 | - |
| | | | | A5302 | KP738581 | KP738541 | KP738501 | OL191323 | PP259890 | - |
| ***Schistura fusinotata*** | Laos | Xekong | Mekong | A5065 | PP279991 | PP315830 | PP280188 | PP280430 | PP259880 | - |
| ***Schistura implicata*** | Vietnam | Lam Dong | No details | GenBank | MG238289 | MG237995 | MG238406 | - | - | - |
| ***Schistura incerta*** | China | No details | No details | GenBank | MK361215 | KP695623 | KP695078 | - | KP695739 | KP694654 |
| ***Schistura* sp. *Lam*** | Vietnam | Nghe An | Lam | A3198 | PP279962 | PP315798 | PP280169 | PP280389 | PP259842 | PP259667 |
| | | | | A3199 | PP279963 | PP315799 | PP280170 | PP280390 | PP259843 | - |
| ***Schistura irregularis*** | Laos | Houaphan | Mekong | CMK25919 | PP279866 | PP315701 | PP280084 | PP280279 | - | PP259635 |
| | | | | _1 | PP279867 | PP315702 | PP280085 | PP280280 | - | - |

| | | | | | | | | | | |
|---|---|---|---|---|---|---|---|---|---|---|
| | | | | CMK25919_2 | | | | | | |
| ***Schistura kaysonei*** | Laos | Bolikhamsay | Mekong | GenBank | NC_031580 | - | - | - | - | - |
| ***Schistura klydonion*** | Laos | Champasak | Mekong | CMK23320_1<br>CMK23320_2 | PP279858<br>PP279859 | PP315693<br>PP315694 | PP280080<br>PP280081 | PP280271<br>PP280272 | PP259980<br>PP259981 | PP259709<br>- |
| ***Schistura kongphengi*** | Vietnam | Thua Thien Hue | Mekong | A2722<br>A3321 | PP279945<br>PP279969 | PP315782<br>PP315805 | PP280155<br>PP280173 | PP280372<br>PP280399 | PP259827<br>PP259851 | -<br>- |
| ***Schistura laterimaculata*** | Thailand | Petchabun | Mekong | A6848<br>A6849 | PP280023<br>PP280024 | PP315856<br>PP315857 | PP280208<br>PP280209 | PP280465<br>PP280466 | PP259922<br>PP259923 | PP259689<br>- |
| ***Schistura latidens*** | China | Yunnan | Mekong | GenBank | MG238266 | MG237973 | MG238383 | - | MG238081 | - |
| ***Schistura latifasciata*** | China | Yunnan | Mekong | GenBank | MG238268 | MG237975 | MG238385 | - | MG238083 | - |
| ***Schistura macrocephala*** | China | Yunnan | Mekong | GenBank | MG238274 | MG237981 | MG238391 | - | MG238088 | - |
| ***Schistura macrotaenia*** | China | Yunnan | Mekong | GenBank | JN837655 | JN837667 | - | - | - | - |
| ***Schistura magnifluvis*** | China | Yunnan | Mekong | GenBank | JN837654 | MG237967 | MG238355 | - | MG238075 | - |
| ***Schistura moeiensis*** | Thailand | Tak | Salween | A4965<br>A4966<br>A11033 | PP279988<br>PP279989<br>PP279883 | PP315826<br>PP315827<br>PP315718 | PP280185<br>PP280186<br>PP280094 | PP280426<br>PP280427<br>PP280296 | PP259876<br>PP259877<br>PP259752 | -<br>-<br>- |
| ***Schistura nicholsi*** | China | Yunnan | Mekong | GenBank | DQ105202 | - | - | - | - | - |
| ***Schistura notasileum*** | China | Yunnan | Mekong | GenBank | OQ945050 | OQ973300 | OQ973298 | - | OQ973302 | - |
| ***Schistura porthos*** | China | Yunnan | Mekong | GenBank | MG238282 | MG237988 | MG238399 | - | - | - |
| ***Schistura reidi*** | Thailand | Mae Hong Son | Salween | A781<br>A817 | PP280032<br>PP280036 | PP315865<br>PP315871 | PP280217<br>PP280220 | PP280488<br>PP280494 | PP259950<br>- | -<br>- |
| ***Schistura rikiki*** | Laos | Xekong | Mekong | A5066<br>A5067 | PP279992<br>PP279993 | PP315831<br>PP315832 | PP280189<br>PP280190 | PP280431<br>PP280432 | PP259881<br>- | PP259682<br>- |
| ***Schistura sertata*** | Laos | Luang Prabang | Mekong | A2531<br>A2531 | PP279936<br>PP279937 | PP315770<br>PP315771 | PP280147<br>PP280148 | PP280361<br>PP280362 | PP259815<br>PP259816 | -<br>- |
| ***Schistura sexcauda*** | Thailand | Chiang Mai | Chao Phraya | A755 | PP280029 | PP315862 | PP280214 | PP280485 | PP259947 | - |
| ***Schistura similis*** | Thailand | Tak | Salween | A5004 | PP279990 | PP315828 | PP280187 | PP280428 | PP259878 | - |
| ***Schistura sokolovi*** | Vietnam | Gia Lei | Song Ba | A3384 | PP279970 | PP315806 | PP280174 | PP280400 | PP259852 | PP259673 |
| ***Schistura susannae*** | Vietnam | Da Nang | Mong Mo | GenBank | MG238288 | MG237994 | MG238405 | - | - | - |
| ***Schistura thanho*** | Vietnam | Bien Dinh | Vinh Thanh | A3310 | PP279968 | PP315804 | - | PP280398 | - | - |

| | | | | | | | | | | |
|---|---|---|---|---|---|---|---|---|---|---|
| ***Schistura waltoni*** | Thailand | Chiang Mai | Chao Phraya | A731 | PP280027 | PP315860 | PP280212 | PP280477 | PP259934 | - |
| | | | | A732 | PP280028 | PP315861 | PP280213 | PP280478 | PP259935 | - |
| ***Schistura xhatensis*** | Laos | Houaphan | Mekong | CMK25920_1 | PP279868 | PP315703 | PP280086 | PP280281 | PP259736 | PP259636 |
| | | | | CMK25920_2 | PP279869 | PP315704 | PP280087 | PP280282 | PP259737 | - |
| ***Schistura yersini*** | Vietnam | Lam Dong | Da Dung | A3206 | PP279964 | PP315800 | PP280171 | PP280391 | PP259844 | PP259668 |
| ***Sectoria atriceps*** | Thailand | Nan | Chao Phraya | A845 | PP280037 | PP315872 | PP280221 | PP280500 | PP259957 | - |
| | | | | A846 | PP280038 | PP315873 | PP280222 | PP280501 | PP259960 | - |
| | | | | A847 | PP280039 | PP315874 | PP280223 | PP280502 | PP259961 | - |
| | | | | A848 | PP280040 | PP315875 | PP280224 | PP280503 | PP259962 | - |
| | | | | A849 | PP280041 | PP315876 | PP280225 | PP280504 | PP259963 | PP259698 |
| ***Sectoria heterognathos*** | Laos | Louang Namtha | Mekong | CMK25980 | PP279872 | PP315705 | PP280089 | PP280283 | PP259740 | - |
| ***Speonectes tiomanensis*** | Malaysia | Pahang | Tioman Island | A1848 | PP279916 | PP315750 | PP280128 | PP280336 | PP259791 | - |
| | | | | A1849 | PP279917 | PP315751 | PP280129 | PP280337 | PP259792 | - |
| ***Tuberoschistura baenzingeri*** | Thailand | Surat Thani | Tapi | A2340 | - | PP315765 | PP280143 | PP280354 | PP259807 | PP259657 |
| | | | | A4485 | - | PP315822 | - | PP280418 | PP259866 | - |
| ***Tuberoschistura cambodgiensis*** | Cambodia | Phnom Penh | Mekong | A7949 | - | PP315866 | PP280218 | PP280489 | PP259951 | PP259696 |
| | | | | A7950 | - | PP315867 | PP280219 | PP280490 | PP259952 | - |
| **SUNDAIC CLADE** | | | | | | | | | | |
| ***Nemacheilus binotatus*** | Thailand | Chiang Mai | Chao Phraya | A6926 | KP738586 | KP738546 | KP738506 | OL191346 | PP259926 | PP259690 |
| | | | | A6927 | KP738587 | KP738547 | PP280210 | PP280469 | PP259927 | - |
| ***Nemacheilus cacao*** | Laos | Bolikhamsay | Mekong | ZRC 62554 | ON720269 | ON720271 | PP280244 | PP280527 | PP259983 | - |
| | | | | ZRC 62553 | ON720270 | ON720272 | PP280245 | PP280528 | - | - |
| ***Nemacheilus* cf. *tuberigum*** | Indonesia | Aceh | Alas | A4536 | PP279986 | PP315823 | - | PP280419 | PP259867 | PP259680 |
| | | | | A4537 | PP279987 | PP315824 | - | PP280420 | PP259868 | - |
| ***Nemacheilus masyae*** | Thailand | Surat Thani | Tapi | A1421 | MW512997 | MW513120 | PP280109 | PP280316 | PP259771 | PP259644 |
| | | | | A1422 | MW512998 | MW513121 | PP280110 | PP280317 | PP259772 | - |
| | Cambodia | Siem Reap | Mekong | A5450 | PP280010 | PP315847 | PP280199 | PP280453 | PP259901 | - |
| ***Nemacheilus ornatus*** | Thailand | Surat Thani | Tapi | A1402 | MW513023 | MW513146 | PP280107 | PP280314 | PP259769 | PP259643 |
| | | | | A1403 | MW513024 | MW513146 | PP280108 | PP280315 | PP259770 | - |
| ***Nemacheilus pallidus*** | Thailand | Nan | Chao Phraya | A1394 | MW513029 | MW513151 | PP280106 | PP280313 | PP259768 | - |
| ***Nemacheilus platyceps*** | Thailand | Chanthaburi | Mekong | A856 | MW513045 | MW513170 | PP280226 | PP280505 | PP259965 | - |
| | | | | A857 | MW513046 | MW513171 | PP280227 | PP280506 | PP259966 | - |

| | | | | | | | | | | |
|---|---|---|---|---|---|---|---|---|---|---|
| ***Nemacheilus saravacensis*** | Malaysia | Sarawak | Sabang | A1632 | MW513056 | MW513181 | PP280114 | PP280321 | PP259775 | - |
| ***Nemacheilus selangoricus*** | Thailand | Nakhon Si Thammarat | Pak Paying | A1437 | MW513062 | MW513186 | PP280111 | PP280318 | - | PP259645 |
| ***Nemacheilus spiniferus*** | Malaysia | Sarawak | Engkaban | A1640 | MW513079 | MW513203 | - | PP280322 | - | PP259647 |
| **BURMESE CLADE** | | | | | | | | | | |
| ***Aborichthys kempi*** | Myanmar | Kachin | Irrawaddy | CMK25538_1 | PP280061 | PP315697 | - | PP280275 | PP259732 | PP259633 |
| | | | | CMK25538_2 | PP280062 | PP315698 | - | PP280276 | PP259733 | - |
| ***Aborichthys* sp.** | India | West Bengal | Brahma-putra | A3972 | PP279977 | PP315814 | PP280177 | PP280407 | PP259857 | - |
| | | | | A3973 | PP279978 | PP315815 | PP280178 | PP280408 | - | - |
| ***Acanthocobitis* cf. *pavonacea*** | Ornamental fish trade | | | A1863 | MK608119 | EF056379 | MK608242 | PP280338 | PP259794 | - |
| | | | | A1864 | MK608120 | MK608146 | MK608243 | PP280339 | PP259795 | - |
| ***Paracanthocobitis epimekes*** | Thailand | Phang Nga | Takua Pa | A2460 | MK608038 | MK608151 | MK608248 | PP280357 | PP259811 | - |
| | Myanmar | Tanintharyi | Tenasserim | CMK 24940 | MK608104 | MK608213 | MK608307 | PP280247 | PP259730 | - |
| ***Paracanthocobitis linypha*** | Myanmar | no details | | A2562 | MK608044 | MK608157 | MK608254 | PP280365 | PP259821 | - |
| | | | | A2563 | MK608045 | MK608158 | MK608255 | PP280366 | - | - |
| ***Paracanthocobitis mackenziei*** | Ornamental | fish trade | | A82 | EF508598 | EF056383 | MK608221 | PP280495 | - | - |
| | | | | A83 | MK608121 | MK608127 | MK608222 | PP280496 | - | - |
| | Nepal | Koshi | Ganges | A3437 | MK608113 | MK608162 | MK608259 | PP280401 | PP259853 | - |
| ***Paracanthocobitis mandalayensis*** | Thailand | Tak | Salween | A11558 | PP279890 | PP315726 | - | PP280304 | PP259759 | - |
| ***Paracanthocobitis phuketensis*** | Myanmar | Tanintharyi | Tenasserim | CMK 28791 | MK608107 | MK608216 | MK608310 | PP280246 | - | - |
| | Thailand | Phatthalung | Phaniat | A5190 | MK608075 | MK608186 | MK608281 | PP280434 | PP259882 | - |
| | Thailand | Trang | Palian | A5175 | MK608071 | MK608182 | MK608277 | PP280433 | - | - |
| | Thailand | Phang Nga | Tam Nang | A9713 | MK608099 | MK608208 | MK608302 | PP280526 | - | - |
| | Thailand | Phang Nga | Takua Pa | A2466 | MK608042 | MK608155 | MK608252 | PP280358 | PP259812 | - |
| ***Paracanthocobitis pictilis*** | Ornamental | fish trade | | A6940 | KP738589 | KP738549 | KP738509 | OL191347 | PP259929 | PP259692 |
| | | | | A6941 | KP738590 | KP738550 | KP738510 | PP280471 | - | - |
| | | | | A10938 | PP279882 | PP315715 | - | PP280293 | PP259749 | - |
| ***Paracanthocobitis* sp. Irrawaddy** | Myanmar | Magway | Irrawaddy | A5774 | MK608078 | MK608189 | MK608284 | PP280456 | PP259905 | - |
| | | | | A6621 | MK608089 | MK608198 | MK608293 | PP280459 | - | - |
| | Myanmar | Ayeyerwady | Irrawaddy | A6573 | MK608086 | MK608196 | MK608291 | PP280458 | PP259912 | - |
| ***Paracanthocobitis* sp. Rakhine** | Myanmar | Rakhine | no details | A5331 | KP738582 | KP738542 | KP738502 | OL191324 | PP259891 | - |
| | | | | A5332 | KP738583 | KP738543 | KP738503 | PP280443 | PP259892 | - |

| | | | | | | | | | | |
|---|---|---|---|---|---|---|---|---|---|---|
| ***Paracanthocobitis* sp. Sittaung** | Myanmar | Mon | Sittaung | A4102 | MK608051 | MK608164 | MK608261 | PP280411 | - | - |
| ***Paracanthocobitis zonalternans*** | Thailand | Tak | Salween | A4942 | MK608065 | MK608176 | MK608271 | PP280425 | - | - |
| ***Schistura* cf. *kohchangensis*** | Thailand | Tak | Salween | A11046 | PP279884 | PP315719 | PP280095 | PP280297 | PP259753 | - |
| ***Schistura kohchangensis*** | Thailand | Chanthaburi | Mekong | A1822<br>A1823 | PP279912<br>PP279913 | PP315746<br>PP315747 | PP280124<br>PP280125 | PP280332<br>PP280333 | PP259787<br>PP259788 | -<br>- |
| ***Schistura savona*** | Ornamental fish trade | | | A7530<br>A7532 | KP738598<br>KP738599 | KP738558<br>KP738559 | KP738518<br>KP738519 | PP280482<br>OL191354 | PP259939<br>PP259940 | PP259695<br>- |
| **SOUTHERN CLADE** | | | | | | | | | | |
| ***Afronemacheilus abyssinicus*** | Ethiopia | Amhara | Nile | A11680 | PP279891 | PP315727 | PP280101 | PP280305 | PP259760 | - |
| ***Mesonoemacheilus guentheri*** | India | Ornamental fish trade | | A2630<br>A2631<br>A6935 | PP279943<br>PP279944 | PP315780<br>PP315781 | PP280153<br>PP280154 | PP280370<br>PP280371<br>PP280470 | PP259825<br>PP259826<br>PP259928 | -<br>-<br>PP259691 |
| ***Mesonoemacheilus herrei*** | India | Tamil Nadu | Periyar | A5526<br>A5527 | PP280011<br>PP280012 | PP315848<br>PP315849 | PP280200<br>PP280201 | PP280454<br>PP280455 | PP259902<br>PP259903 | -<br>- |
| ***Mesonoemacheilus pambarensis*** | India | Tamil Nadu | Pambar | GenBank | MF680101 | - | - | - | - | - |
| ***Mesonoemacheilus petrubanarescui*** | India | Karnataka | Saklshpur | A9197 | PP280055 | PP315890 | PP280241 | PP280520 | PP259979 | - |
| ***Mesonoemacheilus tambaraparniensis*** | India | Tamil Nadu | Tamiraparani | GenBank | MF680096 | - | - | - | - | - |
| ***Mesonoemacheilus triangularis*** | India | Ornamental fish trade | | A944<br>A945 | PP280058<br>PP280059 | PP315891<br>PP315892 | PP280242<br>PP280243 | PP280521<br>PP280522 | PP259984<br>PP259985 | -<br>- |
| ***Mustura bella*** | Laos | Louang Namtha | Mekong | CMK 26052 | OL191242 | OL191491 | OL345558 | OL191373 | PP259743 | - |
| ***Mustura celata*** | Myanmar | Kachin | Irrawaddy | CMK25620_1<br>CMK25620_2 | PP279863<br>PP279864 | PP315699<br>PP315700 | PP280082<br>PP280083 | PP280277<br>PP280278 | PP259734<br>PP259735 | PP259634<br>- |
| ***Mustura geisleri*** | Thailand | Chiang Mai | Chao Phraya | A1238<br>A1239 | OL191158<br>OL191159 | OL191407<br>OL191408 | OL345477<br>OL345478 | OL191270<br>OL191271 | PP259764<br>PP259765 | -<br>- |
| ***Mustura isostigma*** | Laos | Khammouane | Mekong | A1307 | PP279899 | - | - | - | - | - |
| ***Mustura maepaiensis*** | Thailand | Mae Hong Son | Salween | A799<br>A804 | PP280033<br>PP280034 | PP315868<br>PP315869 | -<br>- | PP280491<br>PP280492 | -<br>- | -<br>- |

| | | | | | | | | | | |
|---|---|---|---|---|---|---|---|---|---|---|
| | | | | A805 | PP280035 | PP315870 | - | PP280493 | - | - |
| ***Mustura shanensis*** | Myanmar | Shan | Salween | A6649 | OL191213 | OL191462 | OL345529 | OL191333 | PP259913 | - |
| | | | | A6773 | OL191220 | OL191469 | OL345536 | OL191340 | - | PP259688 |
| | | | | A6774 | OL191221 | OL191470 | OL345537 | OL191341 | PP259920 | - |
| ***Mustura* sp.** | Myanmar | Magway | Irrawaddy | A6712 | PP280019 | PP315852 | PP280204 | PP280461 | PP259917 | - |
| | | | | A6713 | PP280020 | PP315853 | PP280205 | PP280462 | - | - |
| ***Mustura* sp. Salween** | Thailand | Tak | Salween | A1776 | PP279907 | - | - | - | - | - |
| ***'Nemacheilus' corica*** | India | Ornamental fish trade | | A6945 | KP738592 | KP738552 | KP738512 | PP280472 | PP259930 | PP259694 |
| | | | | A6948 | KP738593 | KP738553 | KP738513 | PP280473 | PP259931 | - |
| | | | | A6953 | KP738594 | KP738554 | KP738514 | PP280475 | PP259932 | - |
| ***Nemachilichthys ruppelli*** | India | Ornamental fish trade | | A4341 | KP738573 | KP738533 | KP738493 | OL191311 | PP259864 | PP259678 |
| | | | | A4345 | KP738574 | KP738534 | KP738494 | PP280417 | PP259865 | PP259679 |
| ***Neonoemacheilus labeosus*** | Thailand | Tak | Salween | A1774 | PP279905 | PP315740 | PP280118 | PP280326 | PP259780 | - |
| | | | | A1775 | PP279906 | PP315741 | PP280119 | PP280327 | PP259781 | - |
| ***Neonoemacheilus peguensis*** | Myanmar | Bago | Sittaung | A877 | PP280046 | PP315881 | PP280232 | PP280511 | PP259971 | - |
| ***Neonoemacheilus* sp. Irrawaddy** | Myanmar | Magway | Irrawaddy | A6287 | PP280017 | PP315850 | PP280202 | PP280457 | PP259909 | - |
| ***Neonoemacheilus* sp.** | Myanmar | Magway | Irrawaddy | A6664 | PP280018 | PP315851 | PP280203 | PP280460 | PP259914 | - |
| | | | | A6734 | PP280021 | PP315854 | PP280206 | PP280463 | - | - |
| | | | | A6735 | PP280022 | PP315855 | PP280207 | PP280464 | - | - |
| ***Oxynoemacheilus angorae*** | No details | | | GenBank | NC_031548 | - | - | - | - | - |
| ***Oxynoemacheilus brandtii*** | Armenia | Lori | Kura | A3901 | PP279975 | PP315812 | - | PP280405 | - | - |
| | | | | A3902 | PP279976 | PP315813 | - | PP280406 | - | - |
| ***Oxynoemacheilus bureschi*** | Bulgaria | Blagoevgrad | Struma | A12147 | PP279895 | PP315731 | PP280102 | PP280309 | PP259991 | PP259641 |
| | | | | A12148 | PP279896 | PP315732 | PP280103 | PP280310 | PP259992 | - |
| ***Oxynoemacheilus cilicic*** | Turkey | Adana | Seyhan | A2758 | PP279947 | PP315784 | PP280157 | PP280374 | PP259828 | - |
| ***Oxynoemacheilus chomanicus*** | Iran | Kurdistan | Tigris | GenBank | KT715806 | - | - | - | - | - |
| ***Oxynoemacheilus euphraticus*** | Turkey | Mus | Euphrat | A1003 | PP279876 | PP315708 | PP280090 | PP280286 | - | - |
| ***Oxynoemacheilus frenatus*** | Turkey | Diyarbakir | Tigris | A2102 | PP279926 | PP315759 | PP280137 | PP280348 | - | - |
| | | | | A2103 | PP279927 | PP315760 | PP280138 | PP280349 | - | - |
| ***Oxynoemacheilus galilaeus*** | Syria | Dara | Jordan | A3890 | PP279973 | PP315810 | - | - | - | - |
| | | | | A3891 | PP279974 | PP315811 | PP280176 | - | - | - |

| | | | | | | | | | | |
|---|---|---|---|---|---|---|---|---|---|---|
| ***Oxynoemacheilus gyndes*** | Iraq | Sulaimaniyah | Tigris | GenBank | MH842968 | MH843091 | - | - | - | - |
| ***Oxynoemacheilus hanae*** | Iraq | Sulaimaniyah | Tigris | GenBank | MH842969 | MH843092 | - | - | - | - |
| ***Oxynoemacheilus* veyselorum** | Turkey | Erzurum | Arax | A1045 | PP279877 | PP315709 | - | PP280287 | - | - |
| ***Oxynoemacheilus insignis*** | Israel | West bank | Jordan | A1961<br>A1962 | PP279920<br>PP279921 | PP315754<br>PP315755 | PP280131<br>PP280132 | PP280342<br>PP280343 | PP259796<br>PP259797 | PP259654<br>- |
| ***Oxynoemacheilus kaynaki* 1** | Turkey | Elazig | Euphrat | A1000 | PP279874 | PP315706 | - | PP280284 | - | - |
| ***Oxynoemacheilus kaynaki* 2** | Turkey | Elazig | Euphrat | A1002 | PP279875 | PP315707 | - | PP280285 | - | - |
| ***Oxynoemacheilus kurdistanicus*** | Iran | Kurdistan | Tigris | GenBank | KU180210 | - | - | - | - | - |
| ***Oxynoemacheilus merga*** | Russia | Dagestan | Rubas | A1109<br>A1110 | PP279885<br>PP279886 | PP315720<br>PP315721 | PP280096<br>- | PP280298<br>PP280299 | -<br>- | PP259637<br>- |
| ***Oxynoemacheilus persa*** | Iran | Fars | Kor | A695<br>A697 | PP280025<br>PP280026 | PP315858<br>PP315859 | PP280211<br>- | PP280474<br>PP280476 | PP259933<br>- | -<br>- |
| ***Oxynoemacheilus phasicus*** | Georgia | Imereti | Rioni | A2057 | PP279924 | - | PP280135 | PP280346 | PP259800 | - |
| ***Oxynoemacheilus pindus*** | Albania | Gjirokaster | Vjosa | A12149<br>A12150 | PP279897<br>PP279898 | PP315733<br>PP315734 | PP280104<br>PP280105 | PP280311<br>PP280312 | PP259993<br>PP259994 | PP259642<br>- |
| ***Oxynoemacheilus* sp. Turkey** | Turkey | No details | No details | GenBank | EU015983 | - | - | - | - | - |
| ***Oxynoemacheilus tongiorgii*** | Iran | Fars | Kor | A2757 | PP279946 | PP315783 | PP280156 | PP280373 | - | - |
| ***Oxynoemacheilus zagrosensis*** | Iran | Kurdistan | Tigris | GenBank | KU180203 | - | - | - | - | - |
| ***Oxynoemacheilus zarzianus*** | Iraq | Al-Sulaimaniyah | Tigris | GenBank | KY849795 | - | - | - | - | - |
| ***Paracobitis atrakensis*** | Iran | Khorasane - Shomali | Atrak | GenBank | MG229862 | - | - | - | - | - |
| ***Paracobitis hircanica*** | Iran | Golestan | Gorgan Roud | A2186<br>A2187<br>A2188 | PP279928<br>PP279929<br>PP279930 | PP315761<br>PP315762<br>PP315763 | PP280139<br>PP280140<br>PP280141 | PP280350<br>PP280351<br>PP280352 | PP259802<br>PP259803<br>PP259804 | PP259656<br>-<br>- |
| ***Paracobitis malapterura*** | Iran | Qom | Emamzadeh Abdollah | GenBank | MG229879 | - | - | - | - | - |
| ***Paracobitis molavii*** | Iran | West Azerbaijan | Tigris | GenBank | MG229860 | - | - | - | - | - |

| | | | | | | | | | | |
|---|---|---|---|---|---|---|---|---|---|---|
| ***Paracobitis persa*** | Iran | Fars | Kor | GenBank | MG229866 | - | - | - | - | - |
| ***Paracobitis rhadinaea*** | Iran | Sistan and Baluchistan | Sistan | GenBank | MG229870 | - | - | - | - | - |
| ***Paraschistura cristata*** | Iran | Khorazan Razavi | Hari | A5262 | PP279995 | PP315833 | - | PP280436 | PP259883 | - |
| | | | | A5263 | PP279996 | PP315834 | - | PP280437 | PP259884 | - |
| | | | | A5264 | PP279997 | PP315835 | - | PP280438 | PP259885 | - |
| | | | | A5265 | PP279998 | PP315836 | - | PP280439 | PP259886 | - |
| | | | | A5266 | PP279999 | PP315837 | - | PP280440 | PP259887 | - |
| ***Paraschistura montana*** | India | No details | Ganges | GenBank | FJ711438 | - | - | - | - | - |
| ***Petruichthys brevis*** | Myanmar | Shan | Salween | A4184 | KP738571 | KP738531 | KP738491 | OL191307 | PP259859 | - |
| | | | | A4185 | KP738572 | KP738532 | KP738492 | OL191308 | PP259860 | - |
| | | | | A6503 | OL191210 | OL191459 | OL345526 | OL191330 | PP259910 | - |
| | | | | A6739 | OL191219 | OL191468 | OL345535 | OL191339 | PP259919 | - |
| ***Physoschistura brunneana*** | Myanmar | Shan | Salween | A578 | OL191138 | OL191386 | OL345460 | OL191249 | PP259906 | PP259687 |
| ***Physoschistura* cf. *rivulicola*** | Myanmar | Shan | Salween | A6806 | OL191223 | OL191472 | OL345539 | OL191343 | PP259921 | - |
| ***Physoschistura* cf. *shuangjiangensis*** | China | Yunnan | Mekong | A2999 | OL191173 | OL191423 | OL345492 | OL191288 | - | - |
| ***Physoschistura mango*** | Myanmar | Shan | Salween | A2253 | PP279931 | PP315764 | PP280142 | PP280353 | PP259805 | - |
| ***Physoschistura pseudobrunneana*** | Thailand | Phayao | Chao Phraya | A850 | OL191155 | OL191403 | OL345474 | OL191266 | PP259964 | - |
| | Thailand | Chiang Rai | Mekong | A1356 | OL191163 | OL191412 | OL345482 | OL191275 | PP259766 | - |
| | | | | A1357 | OL191164 | OL191413 | OL345483 | OL191276 | PP259767 | - |
| ***Physoschistura rivulicola*** | Myanmar | Shan | Salween | A6670 | OL191214 | OL191463 | OL345530 | OL191334 | PP259915 | - |
| ***Physoschistura shuangjiangensis*** | China | Yunnan | Mekong | GenBank | MG238284 | MG237990 | MG238401 | - | - | - |
| ***Physoschistura* sp.** | Myanmar | Shan | Salween | A2256 | OL191165 | OL191415 | OL345484 | OL191280 | PP259806 | - |
| | | | | A7545 | KP738600 | KP738560 | KP738520 | OL191355 | PP259943 | - |
| | | | | A7546 | KP738601 | KP738561 | KP738521 | OL191356 | PP259944 | - |
| ***Pteronemacheilus lucidorsum*** | Myanmar | Shan | Irrawaddy | A6695 | OL191215 | OL191464 | OL345531 | OL191335 | PP259916 | - |
| | | | | A8465 | KP738606 | KP738566 | KP738526 | OL191360 | PP259958 | - |
| | | | | A8466 | KP738607 | KP738567 | KP738527 | OL191361 | PP259959 | - |
| ***Pteronemacheilus meridionalis*** | China | Yunnan | Salween | A3122 | PP279958 | PP315795 | - | PP280385 | - | - |
| | Laos | Louang Namtha | Mekong | CMK 26015 | PP279873 | PP315676 | - | PP280250 | - | - |
| | | | | CMK 25928 | PP279870 | PP315675 | - | PP280248 | - | - |

| | | | | | | | | | | |
|---|---|---|---|---|---|---|---|---|---|---|
| ***Pteronemacheilus* sp.** | Myanmar | Shan | Irrawaddy | A5851 | OL191208 | OL191457 | OL345524 | OL191328 | PP259907 | - |
| | | | | A5852 | OL191209 | OL191458 | OL345525 | OL191329 | PP259908 | - |
| ***Sasanidus kermanshahensis*** | Iran | Kermanshah | Tigris | A5261 | PP279994 | - | - | PP280435 | - | - |
| ***Schistura albirostris*** | China | Yunnan | Irrawaddy | GenBank | MG238242 | - | MG238356 | - | - | - |
| ***Schistura ataranensis*** | Myanmar | Ornamental | fish trade | A2560 | MK886975 | PP315774 | MK886884 | PP280364 | PP259820 | - |
| | | | | A5062 | MK886998 | PP315829 | MK886907 | PP280429 | PP259879 | - |
| | Thailand | Kanchanaburi | Ataran | A11005 | MK887031 | PP315716 | MK886939 | PP280294 | PP259750 | - |
| ***Schistura aurantiaca*** | Myanmar | Mon | Ataran | A954 | MK886950 | PP315893 | MK886861 | PP280523 | PP259986 | - |
| | Thailand | Tak | Mae Klong | A9580 | MK887018 | OL191503 | MK886926 | OL191385 | PP259987 | PP259710 |
| | Thailand | Tak | Salween | A9584 | MK887019 | PP315894 | MK886927 | PP280524 | PP259988 | - |
| | | | | A11010 | MK887032 | PP315717 | MK886940 | PP280295 | PP259751 | - |
| ***Schistura balteata*** | Myanmar | Ornamental fish trade | | A2554 | MK886971 | OL191502 | MK886880 | OL191384 | PP259819 | - |
| | | | | A2555 | MK886972 | PP315773 | MK886881 | PP280363 | - | - |
| ***Schistura beavani*** | India | No detail | Ganges | GenBank | GQ478448 | - | - | - | - | - |
| ***Schistura callidora*** | Myanmar | Shan state | Irrawaddy | A3909 | OL191189 | OL191438 | OL345507 | OL191303 | - | - |
| | | | | A3910 | OL191190 | OL191439 | OL345508 | OL191304 | - | - |
| ***Schistura* cf. *nilgiriensis*** | India | Karnataka | Netravanthi | A9199 | PP280057 | - | - | - | - | - |
| ***Schistura* cf. *poculi* 1** | Thailand | Tak | Mae Klong | A4202 | OL191193 | OL191442 | OL345511 | OL191309 | - | PP259675 |
| | | | | A4203 | OL191194 | OL191443 | OL345512 | OL191310 | - | - |
| ***Schistura* cf. *poculi* 2** | China | Yunnan | Salween | A2914 | OL191172 | OL191422 | OL345491 | OL191287 | PP259829 | - |
| ***Schistura* cf. *scaturigina*** | India | West Bengal | Rydak I River | A3925 | OL191191 | OL191440 | OL345509 | OL191305 | PP259856 | - |
| | | | | A3926 | OL191192 | OL191441 | OL345510 | OL191306 | - | - |
| ***Schistura* cf. *sijuensis*** | India | Ornamental fish trade | | A3698 | OL191184 | OL191434 | OL345503 | OL191299 | - | - |
| | | | | A3699 | OL191185 | OL191435 | OL345504 | OL191300 | - | - |
| | | | | A11327 | OL191248 | OL191499 | OL345566 | OL191381 | - | - |
| ***Schistura* cf. *vinciguerrae*** | Myanmar | Rakhine | Irrawaddy | A5557 | OL191205 | OL191454 | OL345521 | OL191325 | PP259904 | - |
| | Myanmar | Magway | Irrawaddy | A6564 | OL191211 | OL191460 | OL345527 | OL191331 | PP259911 | - |
| | | | | A6736 | OL191216 | OL191465 | OL345532 | OL191336 | PP259918 | - |
| ***Schistura cincticauda*** | Thailand | Tak | Salween | A8312 | MK887016 | - | MK886924 | PP280497 | PP259953 | - |
| | | | | A8313 | MK887017 | - | MK886925 | PP280498 | PP259954 | - |
| ***Schistura conirostris*** | China | Yunnan | Mekong | GenBank | MG238247 | MG237953 | MG238362 | - | MG238061 | - |
| ***Schistura crabro*** | Laos | Bolikhamsai | Mekong | CMK 24559 | OL191231 | OL191480 | OL345547 | OL191362 | PP259982 | - |
| ***Schistura crocotula*** | Thailand | Prachuap Khiri Khan | Bang Saphan | A9591 | MK887021 | PP315895 | MK886928 | PP280525 | PP259989 | - |
| | | | | A10513 | MK887024 | PP315713 | MK886931 | PP280290 | PP259746 | - |

| | | | | | | | | | | |
|---|---|---|---|---|---|---|---|---|---|---|
| ***Schistura denisoni*** | India | Tamil Nadu | Vaigai | A5531 | PP280015 | - | - | - | - | - |
| | | | | A5532 | PP280016 | - | - | - | - | - |
| ***Schistura devdedi*** | Nepal | No detail | Ganges | A3445 | PP279972 | PP315808 | PP280175 | PP280403 | - | - |
| | India | Ornamental fish trade | | A7541 | KP738608 | KP738568 | KP738528 | PP280483 | PP259941 | - |
| | | | | A7542 | KP738609 | KP738569 | KP738529 | PP280484 | PP259942 | - |
| ***Schistura disparizona*** | China | Yunnan | Salween | GenBank | MG238252 | MG238368 | MG237958 | - | MG238065 | - |
| ***Schistura hartli*** | Thailand | Surat Thani | Tapi | A3690 | MK886978 | PP315809 | MK886887 | PP280404 | PP259855 | - |
| ***Schistura hoai*** | Laos | Houaphan | Mekong | CMK25918_1 | OL191238 | OL191487 | OL345554 | OL191369 | - | - |
| | | | | CMK25918_2 | OL191239 | OL191488 | OL345555 | OL191370 | - | - |
| ***Schistura hypsiura*** | Myanmar | Ornamental fish trade | | A6922 | KP738584 | KP738544 | KP738504 | PP280467 | PP259924 | - |
| | | | | A6925 | KP738585 | KP738545 | KP738505 | PP280468 | PP259925 | - |
| ***Schistura indawgyiana*** | Myanmar | Kachin | Irrawaddy | CMK 25633 | PP279865 | - | - | - | - | - |
| ***Schistura jarutanini*** | Thailand | Kanchanaburi | Mae Klong | GenBank | NC_031584 | - | - | - | - | - |
| ***Schistura kloetzliae*** | Laos | Louang Namtha | Mekong | CMK25994_1 | OL191240 | OL191489 | OL345556 | OL191371 | PP259741 | - |
| | | | | CMK25994_2 | OL191241 | OL191490 | OL345557 | OL191372 | PP259742 | - |
| | | | | GenBank | MG238237 | MG237945 | MG238351 | - | - | - |
| ***Schistura kuehnei*** | Thailand | Surat Thani | Tapi | A4672 | MK886991 | PP315825 | MK886900 | PP280422 | PP259871 | |
| | | | | A11261 | MK887040 | PP315724 | PP280099 | PP280302 | PP259756 | |
| ***Schistura longa*** | China | Yunnan | Salween | GenBank | MG238272 | MG237979 | MG238389 | - | MG238086 | - |
| ***Schistura mahnerti*** | Thailand | Mae Hong Son | Salween | A778 | PP280030 | PP315863 | PP280215 | PP280486 | PP259948 | - |
| | | | | A779 | PP280031 | PP315864 | PP280216 | PP280487 | PP259949 | - |
| ***Schistura malaisei*** | Myanmar | Kachin | Irrawaddy | CMK25506_1 | PP279860 | - | - | - | - | - |
| | | | | CMK25506_2 | PP279861 | PP315696 | - | - | - | - |
| ***Schistura mukambbikaensis*** | India | Karnataka | Netravathi | A9198 | PP280056 | - | - | - | - | - |
| ***Schistura myaekanbawensis*** | Myanmar | Tanintharyi | Tenasserim | CMK24993 | MK887022 | PP315695 | MK886929 | PP280273 | PP259731 | - |
| ***Schistura nilgiriensis*** | India | Ornamental fish trade | | A2626 | PP279942 | PP315779 | PP280152 | PP280369 | PP259824 | - |

| | | | | | | | | | | |
|---|---|---|---|---|---|---|---|---|---|---|
| ***Schistura notostigma*** | Sri Lanka | Ornamental fish trade | | A7519<br>A7520<br>A7521 | KP738595<br>KP738596<br>KP738597 | KP738555<br>KP738556<br>KP738557 | KP738515<br>KP738516<br>KP738517 | PP280479<br>PP280480<br>PP280481 | PP259936<br>PP259937<br>PP259938 | -<br>-<br>- |
| ***Schistura nubigena*** | Myanmar | Kachin | Irrawaddy | CMK 25509 | PP279862 | - | - | PP280274 | PP259990 | - |
| ***Schistura obliquofascia*** | India | No details | No details | GenBank | HM636831 | - | - | - | - | - |
| ***Schistura paucicincta*** | Thailand | Tak | Salween | A4946<br>A4947 | OL191203<br>OL191204 | OL191452<br>OL191453 | OL345519<br>OL345520 | OL191321<br>OL191322 | PP259874<br>PP259875 | -<br>- |
| ***Schistura polytaenia*** | China | Yunnan | Irrawaddy | GenBank | MG238280 | MG237986 | MG238397 | - | MG238092 | - |
| ***Schistura pridii*** | Thailand | Ornamental fish trade | | A7548<br>A7549 | KP738602<br>KP738603 | KP738562<br>KP738563 | KP738522<br>KP738523 | OL191357<br>OL191358 | PP259945<br>PP259946 | -<br>- |
| ***Schistura reticulofasciata*** | India | Assam | Brahmaputra | GenBank | KY379150 | - | - | - | - | - |
| ***Schistura robertsi*** | Thailand | Phang Nga | Tam Nang | A2424 | MK886959 | OL191501 | MK886869 | OL191383 | PP259810 | - |
| ***Schistura rupecula*** | Nepal | No detail | Ganges | A3443 | PP279971 | PP315807 | - | PP280402 | PP259854 | - |
| ***Schistura semiarmatus*** | India | Tamil Nadu | Vaigai | A5529<br>A5530 | PP280013<br>PP280014 | -<br>- | -<br>- | -<br>- | -<br>- | -<br>- |
| ***Schistura sikmaensis*** | China | Yunnan | Irrawaddy | GenBank | JF340405 | JF340413 | - | - | - | - |
| ***Schistura* sp. *Myanmar*** | Myanmar | Ornamental fish trade | | A2569<br>A2570 | PP279938<br>PP279939 | PP315775<br>PP315776 | -<br>- | -<br>- | -<br>- | -<br>- |
| ***Schistura thavonei*** | Laos | Louang Namtha | Mekong | CMK 26066<br>CMK25944_1 | OL191243<br>OL191244 | OL191492<br>OL191493 | OL345559<br>OL345560 | OL191374<br>OL191375 | PP259744<br>PP259739 | -<br>- |
| ***Schistura tirapensis*** | India | Ornamental fish trade | | A3703<br>A3704 | OL191187<br>OL191188 | OL191436<br>OL191437 | OL345505<br>OL345506 | OL191301<br>OL191302 | -<br>- | -<br>- |
| ***Schistura udomritthiruji*** | Thailand | Ranong | Kapoe | A2546<br>A2547 | OL191168<br>MK886969 | OL191418<br>PP315772 | OL345487<br>PP280149 | OL191283<br>- | PP259817<br>PP259818 | PP259661<br>- |
| ***Schistura yingjiangensis*** | China | Yunnan | Irrawaddy | GenBank | MG238294 | MG237999 | MG238411 | - | MG238103 | - |
| ***Seminemacheilus ispartensis*** | Turkey | Isparta | Egidir | A4833<br>A4834 | KP738577<br>KP738578 | KP738537<br>KP738538 | KP738497<br>KP738498 | PP280423<br>PP280424 | PP259872<br>PP259873 | -<br>- |
| ***Turcinoemacheilus ekmekciae*** | Turkey | Diyarbakir | Tigris | A2089 | PP279925 | PP315758 | PP280136 | PP280347 | PP259801 | - |
| ***Turcinoemacheilus* sp. 1** | Iran | Kurdistan | Choman | GenBank | KT861416 | - | - | - | - | - |
| ***Turcinoemacheilus* sp. 2** | Iran | Khuzestan | Karoon | GenBank | GQ338827 | - | - | - | - | - |

| | | | | | | | | | | |
|---|---|---|---|---|---|---|---|---|---|---|
| **OUTGROUP** | | | | | | | | | | |
| **Catostomidae** | | | | | | | | | | |
| ***Cycleptus elongatus*** | | | | GenBank | NC_031634 14392-15532 | EU409613 | EU409671 | - | EU409639 | EU409767 |
| ***Catostomus commersonii*** | | | | GenBank | JX488781 | EU409612 | EU409670 | FJ918841 | EU409638 | EU409766 |
| ***Hypentelium nigricans*** | | | | GenBank | AF454909 | EU711134 | JX470004 | JX190418 | FJ197033 | - |
| **Gyrinocheilidae** | | | | | | | | | | |
| ***Gyrinocheilus aymonieri*** | | | | GenBank | NC_008672 14393-15533 | EU292682 | FJ197122 | - | FJ197071 | EU409791 |
| ***Gyrinocheilus pennocki*** | | | | GenBank | NC_031544 14395-15535 | FJ650415 | FJ650486 | - | FJ650474 | FJ650461 |
| **Botiidae** | | | | | | | | | | |
| ***Leptobotia pellegrini*** | | | | GenBank | NC_031602 14385-15525 | EU292683 | EU409672 | - | EU409640 | EU409768 |
| ***Botia dario*** | | | | GenBank | KU517084 | KU517026 | MF681756 | - | EU409641 | - |
| ***Yasuhikotakia morleti*** | | | | GenBank | NC_031600 14377-15517 | FJ650412 | FJ650483 | - | FJ650471 | FJ650457 |
| ***Syncrossus beauforti*** | | | | GenBank | NC_031546 14387-15527 | FJ650411 | FJ650482 | - | FJ650470 | FJ650456 |
| **Vaillantellidae** | | | | | | | | | | |
| ***Vaillantella maassi*** | | | | GenBank | NC_008680 14378-15518 | EU711132 | FJ197080 | - | FJ197031 | FJ650469 |
| **Cobitidae** | | | | | | | | | | |
| ***Pangio oblonga*** | | | | GenBank | NC_031592 14386-15526 | EU711141 | FJ197091 | - | FJ197041 | FJ650459 |
| ***Canthophrys gongota*** | | | | GenBank | NC_031576 14380-15516 | FJ650414 | FJ650485 | - | FJ650473 | FJ650460 |
| ***Cobitis takatsuensis*** | | | | GenBank | NC_015306 14445-15585 | EU409616 | EU409675 | - | EU409643 | EU409771 |
| ***Cobitis lutheri*** | | | | GenBank | JN858887 | EF508614 | KM818238 | KM818242 | KM583634 | - |
| ***Cobitis taenia*** | Germany | Lower Saxonia | Weser | A1860 | EF508508 | EF056334 | MK608315 | OL191279 | PP259793 | PP259653 |
| ***Cobitis multifasciata*** | | | | GenBank | NC_027166 14376-15516 | EU409615 | EU409674 | - | EU409642 | EU409770 |

| | | | | | | | |
|---|---|---|---|---|---|---|---|
| ***Cobitis tetralineata*** | GenBank | KF661673 | OK661770 | - | OK661564 | - | - |
| **Ellopostomatidae** | | | | | | | |
| ***Ellopostoma mystax*** | GenBank | NC_031642 14377-15517 | FJ650417 | FJ650489 | - | FJ650477 | FJ650464 |
| **Balitoridae** | | | | | | | |
| ***Homaloptera parclitella*** | GenBank | NC_031634 14392-15532 | EU409610 | EU409668 | - | EU409636 | EU409764 |
| **Gastromyzonidae** | | | | | | | |
| ***Sewellia lineolata*** | GenBank | NC_015534 14375-515 | EU409609 | EU409667 | - | EU409635 | EU409763 |
| **Cyprinidae** | | | | | | | |
| ***Enteromius callipterus*** | GenBank | KP712230 | FJ531247 | FJ531365 | - | FJ531345 | FJ531317 |

**Table S2.**

List of primers used in the present study for amplification and/or sequencing

| Locus | Primer name | Primer sequence (5' - 3') | Reference |
|---|---|---|---|
| Cyt *b* | Glu-L.Ca14337–14359 | GAA GAA CCA CCG TTG TTA TTC AA | Šlechtová et al., 2006 |
| | Thr-H.Ca15568–15548 | *ACC TCC RAT CTY CGG ATT ACA* | Šlechtová et al., 2006 |
| | CB-L.Ca14975–14994 | CAC GAR ACR GGR TCN AAY AA | Šlechtová et al., 2006 |
| | CB-H.Ca15057–15035 | *TCT TTR TAT GAG AAR TAN GGG TG* | Šlechtová et al., 2006 |
| IRBP 2 | 101F | TCM TGG ACA AYT ACT GCT CAC C | Chen et al., 2008 |
| | 109F | AAC TAC TGC TCR CCA GAA AAR C | Chen et al., 2008 |
| | 1001R | *GGA AAT GCA TAG TTG TCT GCA A* | Chen et al., 2008 |
| | 1162R | *TGG TGG WCT TYA GGC ACT TGT* | Chen et al., 2008 |
| RAG 1 | RAG-1F | AGC TGT AGT CAG TAY CAC AAR ATG | Grande et al., 2004 |
| | RAG-RV1 | *TCC TGR AAG ATY TTG TAG AA* | Šlechtová et al., 2007 |
| MYH 6 | myh6-F507 | GGA GAA TCA RTC KGT GCT CAT CA | Li et al., 2007 |
| | myh6-R1322 | *CTC ACC ACC ATC CAG TTG AAC AT* | Li et al., 2007 |
| RH 1 | RH-1F | CAT ACG AAT ATC CCC AGT ACT ACC | Liu et al., 2012 |
| | RH-28F | TAC GTG CCT ATG TCC AAY GC | Chen et al. 2008 |
| | RH-139F | CNT ATG AAT AYC CTC AGT ACT ACC | Chen et al. 2003 |
| | RH-233F | ATA TGC CTG CCT GGC YGC TTA C | Chen et al. 2008 |
| | RH-1R | *GCT TGT TCA TGC AGA TGT AGA TGC* | Liu et al., 2012 |
| | RH-1039R | *TGC TTG TTC ATG CAG ATG TAG A* | Chen et al. 2003 |
| EGR 3 | E3-161F | AAT ATC ATG GAC YTG GGN ATG G | Chen et al. 2008 |
| | E3-1136R | *GGY TTC TTG TCC TTC TGT TTS AG* | Chen et al. 2008 |

Chen, W.-J., Miya, M., Saitoh, K., Mayden, R.L., 2008. Phylogenetic utility of two existing and four novel nuclear gene loci in reconstructing Tree of Life of ray-finned fishes: The order Cypriniformes (Ostariophysi) as a case study. Gene 423, 125–134.

Grande, T., Laten, H., Lopez, J.A., 2004. Phylogenetic relationships of extant esocid species (Teleostei: Salmoniformes) based on morphological and molecular characters. Copeia 743–757.

Li, C., Orti, G., Zhang, G., Lu, G. 2007. A Practical Approach to Phylogenomics: The Phylogeny of Ray-Finned Fish (Actinopterygii) as a Case Study. BMC Evol. Biol. 7, 44.

Liu, S.-Q.; Mayden, R.L.; Zhang, J.-B.; Yu, D.; Tang, Q.-Y.; Deng, X.; Liu, H.-Z. Phylogenetic relationships of the Cobitoidea (Teleostei: Cypriniformes) inferred from mitochondrial and nuclear genes with analyses of gene evolution. *Gene* **2012**, *508*, 60– 72.

Šlechtová, V., Bohlen, J., Freyhof, J., Ráb, P., 2006. Molecular phylogeny of the Southeast Asian freshwater fish family Botiidae (Teleostei: Cobitoidea) and the origin of polyploidy in their evolution. Mol. Phylogenet. Evol. 39, 529–541.

Šlechtová, V., Bohlen, J. Tan, H.-H. 2007. Families of Cobitoidea (Teleostei; Cypriniformes) as revealed from nuclear genetic data and the position of the mysterious genera Barbucca, Psilorhynchus, Serpenticobitis and Vaillantella. Mol. Phylogenet. Evol. 44, 1358-65.

**Table S3.**

Alignment attributes and best-fit models. Lengths of alignments, numbers of variable (VP) and parsimony informative (PI) positions and models estimated for all partitions. BEAST and MrBayes models were calculated in Partition Finder 2 (PF2, Lanfear et al., 2016) implemented in PhyloSuite v1.2.2 (Zhang et al., 2020) under AICc criterion, with greedy algorithm (Lanfear et al., 2016) and branch lengths linked. For ML trees the models and partitioning schemes were estimated under BIC by ModelFinder (Kalyaanamoorthy et al. 2017) implemented in IQ tree. The values and models were calculated for both (A) full as well as (B) reduced dataset. Table (C) provides an overview of data attributes for the ingroup dataset only.

**A: full dataset**

| locus | | EGR3 | IRBP 2 | MYH6 | RAG1 | RH | Cytb |
|---|---|---|---|---|---|---|---|
| length (bp) | | 876 | 831 | 777 | 950 | 844 | 1122 |
| Variable positions (VP) | | 341 | 559 | 331 | 503 | 385 | 680 |
| % VP | | 38.93 | 67.27 | 42.60 | 52.95 | 45.62 | 60.61 |
| Pars. Informative (PI) | | 268 | 483 | 299 | 453 | 333 | 615 |
| % PI | | 30.59 | 58.12 | 38.48 | 47.68 | 39.45 | 54.81 |
| | partition | | | | | | |
| MrBayes models (PF2, AICc) | gene | GTR+I+G | SYM+I+G | GTR+I+G | SYM+I+G | GTR+I+G | GTR+I+G |
| | 1st c.p. | GTR+I+G | GTR+I+G | GTR+I+G | GTR+I+G | SYM+I+G | SYM+I+G |
| | 2nd c.p. | GTR+I+G | GTR+I+G | GTR+I+G | GTR+I+G | GTR+I+G | GTR+I+G |
| | 3rd c.p. | GTR+I+G | SYM+I+G | SYM+G | SYM+I+G | GTR+G | GTR+G |
| IQ-Tree models (ModelFinder, BIC) | gene | TPM2u+F+I+G4 | TIM2e+I+G4 | TIM2e+I+G4 | TIM2e+I+G4 | TIM2e+I+G4 | TVM+F+I+G4 |
| | 1st c.p. | TIM2e+I+G4 | TIM3+F+R3 | TN+F+R3 | TIM2e+R3 | TIM2e+R3 | TIM2e+I+G4 |
| | 2nd c.p. | TIM3e+R2 | TVM+F+G4 | TIM2+F+R3 | TVMe+I+G4 | TPM2+F+R3 | TVM+F+R4 |
| | 3rd c.p. | GTR+F+G4 | TIM2e+R3 | TIM2e+R4 | TVMe+G4 | TIM2+F+G4 | TIM2+F+ASC+R5 |
| BEAST models (PF2, AICc) | gene | GTR+I+G+X | GTR+I+G+X | GTR+I+G+X | GTR+I+G+X | GTR+I+G+X | GTR+I+G+X |
| | 1st c.p. | N/E | N/E | N/E | N/E | N/E | N/E |
| | 2nd c.p. | N/E | N/E | N/E | N/E | N/E | N/E |

| | 3rd c.p. | N/E | N/E | N/E | N/E | N/E | N/E |
|---|---|---|---|---|---|---|---|

**Table S3 continuation**

**B: reduced dataset**

| locus | | EGR3 | IRBP 2 | MYH6 | RAG1 | RH | Cytb |
|---|---|---|---|---|---|---|---|
| alignment bp | | 876 | 831 | 777 | 950 | 844 | 1122 |
| var.pos. | | 341 | 546 | 327 | 500 | 376 | 681 |
| % V | | 38.93 | 65.70 | 42.08 | 52.63 | 44.55 | 60.70 |
| Pars. Inf. | | 268 | 436 | 284 | 427 | 312 | 579 |
| % PI | | 30.59 | 52.47 | 36.55 | 44.95 | 36.97 | 51.60 |
| | partition | | | | | | |
| MrBayes models (PF2, AICc) | gene | GTR+I+G | SYM+I+G | GTR+I+G | SYM+I+G | GTR+I+G | GTR+I+G |
| | 1st c.p. | GTR+I+G | GTR+I+G | GTR+I+G | SYM+I+G | SYM+I+G | SYM+I+G |
| | 2nd c.p. | GTR+I+G | GTR+I+G | GTR+I+G | GTR+I+G | GTR+I+G | GTR+I+G |
| | 3rd c.p. | GTR+I | SYM+I+G | SYM+G | SYM+I+G | GTR+G | GTR+G |
| IQ-Tree models (ModelFinder, BIC) | gene | HKY+F+I+G4 | TIM2e+I+G4 | TN+F+I+G4 | TIM2e+I+G4 | TPM2+F+I+G4 | GTR+F+I+G4 |
| | 1st c.p. | TIM2+F+I+G4 | TIM3+F+G4 | K2P+I+G4 | TIM2e+I+G4 | TNe+I+G4 | TIM2e+I+G4 |
| | 2nd c.p. | TIM2+F+I+G4 | TVM+F+G4 | TIM2+F+I+G4 | TIM2e+I+G4 | TVM+F+I+G4 | TVM+F+G4 |
| | 3rd c.p. | K2P+G4 | TIM2e+G4 | TIM2e+G4 | TVMe+I+G4 | TIM2e+G4 | TIM2+F+ASC+G4 |
| BEAST models (PF2, AICc) | gene | GTR+I+G+X | GTR+I+G+X | GTR+I+G+X | GTR+I+G+X | GTR+I+G+X | GTR+I+G+X |
| | 1st c.p. | HKY+I+G+X | TRN+I+G+X | TRN+I+G+X | GTR+I+G+X | GTR+I+G+X | GTR+I+G+X |
| | 2nd c.p. | GTR+I+G+X | GTR+G+X | GTR+I+G+X | GTR+I+G+X | GTR+I+G+X | GTR+I+G+X |
| | 3rd c.p. | GTR+G+X | GTR+I+G+X | GTR+G+X | GTR+I+G+X | GTR+G+X | GTR+G+X |

**Table S3 continuation**

**C: full dataset only ingroup**

| Locus | length (bp) | VP | % VP | PI | % PI |
|---|---|---|---|---|---|
| EGR3 | 876 | 286 | 32.65 | 207 | 23.63 |
| IRBP 2 | 831 | 524 | 63.06 | 443 | 53.31 |
| MYH6 | 777 | 313 | 40.28 | 286 | 36.81 |
| RAG1 | 950 | 463 | 48.74 | 425 | 44.74 |
| RH | 844 | 352 | 41.71 | 307 | 36.37 |
| Cytb | 1122 | 668 | 59.54 | 610 | 54.37 |

Lanfear, R., Calcott, B., Ho, S. Y., & Guindon, S. (2012). PartitionFinder: combined selection of partitioning schemes and substitution models for phylogenetic analyses. Molecular biology and evolution, 29(6), 1695-1701.

Lanfear, R., Frandsen, P. B., Wright, A. M., Senfeld, T., Calcott, B. (2016) PartitionFinder 2: new methods for selecting partitioned models of evolution for molecular and morphological phylogenetic analyses. Molecular biology and evolution. DOI: dx.doi.org/10.1093/molbev/msw260

Kalyaanamoorthy S, Minh BQ, Wong TKF, von Haeseler A, Jermiin LS. 2017. ModelFinder: fast model selection for accurate phylogenetic estimates. Nat Methods. 14(6):587–589.

Zhang, D.; Gao, F.; Jakovlić , I.; Zou, H.; Zhang, J.; Li, W.X.; Wang, G.T. PhyloSuite: An integrated and scalable desktop platform for streamlined molecular sequence data management and evolutionary phylogenetics studies. Mol. Ecol. Resour. 2020, 20,348–355

**Table S4.**

Comparison of biogeographic models in RASP. The last column shows the p-values of the Likelihood Ratio Test. Based on AICc weight (AICc wt), the DEC+J model is recommended as the best fit for our dataset, supported by low p-values indicating the significant influence of the J factor on model likelihood.

| Model | LnL | numparams | d | e | j | AICc | AICc_wt | LRT p-val |
|---|---|---|---|---|---|---|---|---|
| DEC | -369.3 | 2 | 0.0029 | 0.047 | 0 | 742.6 | 2.0e-26 | 1.2e-27 |
| DEC+J | -309.9 | 3 | 1.0e-12 | 0.017 | 0.013 | 625.8 | 0.45 | |
| DIVALIKE | -367.2 | 2 | 0.0031 | 0.046 | 0 | 738.5 | 1.5e-25 | 3.3e-26 |
| DIVALIKE+J | -311.2 | 3 | 1.0e-12 | 0.044 | 0.014 | 628.4 | 0.13 | |
| BAYAREALIKE | -337.9 | 2 | 0.0044 | 0.22 | 0 | 679.8 | 8.7e-13 | 7.8e-14 |
| BAYAREALIKE+J | -310 | 3 | 0.0002 | 0.15 | 0.012 | 626 | 0.42 | |
